# PICNIC accurately predicts condensate-forming proteins regardless of their structural disorder across organisms

**DOI:** 10.1101/2023.06.01.543229

**Authors:** Anna Hadarovich, Hari Raj Singh, Soumyadeep Ghosh, Nadia Rostam, Anthony A. Hyman, Agnes Toth-Petroczy

## Abstract

Biomolecular condensates are membraneless organelles that can concentrate hundreds of different proteins to operate essential biological functions. However, accurate identification of their components remains challenging and biased towards proteins with high structural disorder content with focus on self-phase separating (driver) proteins. Here, we present a machine learning algorithm, PICNIC (Proteins Involved in CoNdensates In Cells) to classify proteins involved in biomolecular condensates regardless of their role in condensate formation. PICNIC successfully predicts condensate members by identifying amino acid patterns in the protein sequence and structure in addition to the intrinsic disorder and outperforms previous methods. We performed extensive experimental validation *in cellulo* and demonstrated that PICNIC accurately predicts 21 out of 24 condensate-forming proteins regardless of their structural disorder content. Even though increasing disorder content was associated with organismal complexity, we found no correlation between predicted condensate proteome content and disorder content across organisms. Overall, we applied a novel machine learning classifier to interrogate condensate components at single protein and whole-proteome levels across the tree of life (picnic.cd-code.org).

## Introduction

Biomolecular condensates are membrane-less organelles that can selectively concentrate biomolecules^1^ and are typically non-stochiometric assemblies of thousands of protein molecules and nucleic acids. The role of condensates has been implicated in several fundamental biochemical processes in physiology and disease^2–4^. Functions exerted by condensates include: i) sequestering molecules and shutting down translation of specific mRNAs^5^; ii) buffering concentration of proteins^6^; iii) reservoir of proteins for fast assembly and disassembly of large complexes, such as the nuclear envelope^7^.

Proteins can have two major roles in a condensate: drivers (scaffolds) or clients^8,9^. Drivers can induce the formation of condensates and are essential members of condensates. For example, knock-out of the driver protein can lead to disassembly of a condensate. While a client is recruited to a condensate often via an interaction with a driver protein, and it is neither necessary nor sufficient in driving the condensate formation. Drivers often self-phase separate *in vitro*, nevertheless self-phase separation does not guarantee *in vivo* driver functionality. For most condensate-forming proteins the client or driver status is unknown, therefore we refer to them as condensate members.

While many proteins can phase-separate in the test-tube and form liquid-like condensates *in vitro*, studying if they also form condensates *in viv*o is more challenging. Individual proteins can be labeled using fluorescent tags and imaged for testing droplet formation which exhibit liquid-like properties such as fusion, Oswald ripening, fast dynamics in fluorescent recovery after photobleaching (FRAP)^10,11^ assay. However, condensates may contain hundreds and even thousands of different proteins. Systematic detection of condensate proteomes is limited to a few mass spectrometry and proximity labeling studies: purified nucleoli^12,13^, P-bodies^14^ and stress granules^15,16^ were subjected to mass spectrometry analysis for enrichment. Systematically and experimentally testing which proteins are members of condensates remains a bottleneck in the field. Therefore, computational methods can facilitate the process of characterizing proteins involved in biomolecular condensates at proteome-scale.

The main limitation of computational method development is related to the sparse experimental data of verified condensate-forming proteins. The first models were proposed based on properties of a few protein families, such as CatGranule^17^ and PScore^18,19^. While these methods aided the discovery of novel condensate-forming proteins with similar properties, they do not generalize well^20^.

In recent years, several data driven and machine learning based liquid-liquid phase separation (LLPS) predictors have been developed^21–24^ making use of the experimental data aggregated in four LLPS databases^25–28^. While most predictors focused on identifying self-phase separating proteins (i.e. form *in vitro* condensates, that are also often drivers)^22–24^ and thus trained on *in vitro* data, a recent metapredictor combined scores of previous methods and microscopy data to identify all condensate members (both drivers and clients)^29^. These methods have excellent performance compared to the first generation of predictors, nevertheless they have several shortcomings. For example a sub-optimal or biased definition of the negative dataset based on a priori assumptions about driving features, such as disorder being the main determinant, and accordingly using structured proteins from PDB as negative data^23^.

Here, we focused on predicting proteins involved in biomolecular condensates instead of proteins involved in *in vitro* self-phase separation per se. We developed a new machine learning model called PICNIC (Proteins Involved in CoNdensates In Cells). As amino acid composition bias and patterning of charges were shown to impact the ability of proteins to form condensates^30–32^, we developed novel features that represent short and long range co-occurrences of amino acids in the protein sequence and structure (AlphaFold2 models), as well as used features such as sequence complexity^23^, disorder score^33^ that were previously shown to be successful in identifying drivers. We addressed the short-comings of previous methods by using a larger and highly curated and non-redundant dataset of *in vivo* condensates (derived from CD-CODE)^34^, and by defining the non-condensate forming, negative dataset, based on a protein-protein interaction network.

Experimental validation of 24 proteins spanning a wide-range structural disorder confirmed that 18 of them localize to condensates with high confidence, while 3 form condensates with low confidence. Further, 8 of them co-localize with known biomolecular condensates. Thus, our experimental validation suggests an ∼88% success rate in identifying condensate forming proteins. Moreover, we trained another model with extended set of features which include Gene Ontology terms (PICNIC_GO_), which provides useful insights about specific protein functions that are enriched in proteins of biomolecular condensate, such as RNA-binding.

Although PICNIC was trained on the richest human data, it generalizes well to other organisms tested. Proteome-wide predictions by PICNIC estimate that ∼40% of proteins partition into condensates across different organisms, from bacteria to humans, with no apparent correlation with organismal complexity or disordered protein content.

## Results

### Defining condensate-forming proteins

In order to develop a model to identify condensate-forming proteins, we assembled a ground truth dataset for *H. sapiens*, that has the most experimentally studied condensates of all organisms to date. Since we aimed at developing a binary classifier, we considered two classes of proteins: 1) proteins involved in condensates (positive dataset) and proteins not involved in condensates (negative dataset) (**Figure 1a**). The positive dataset was constructed from a semi-manually curated dataset of biomolecular condensates and their respective proteins, called CD-CODE (CrowDsourcing COndensate Database and Encyclopedia), developed by our labs^34^. CD-CODE compiles information from primary literature and from four widely used databases of LLPS proteins ^25–28^.

**Figure 1.**
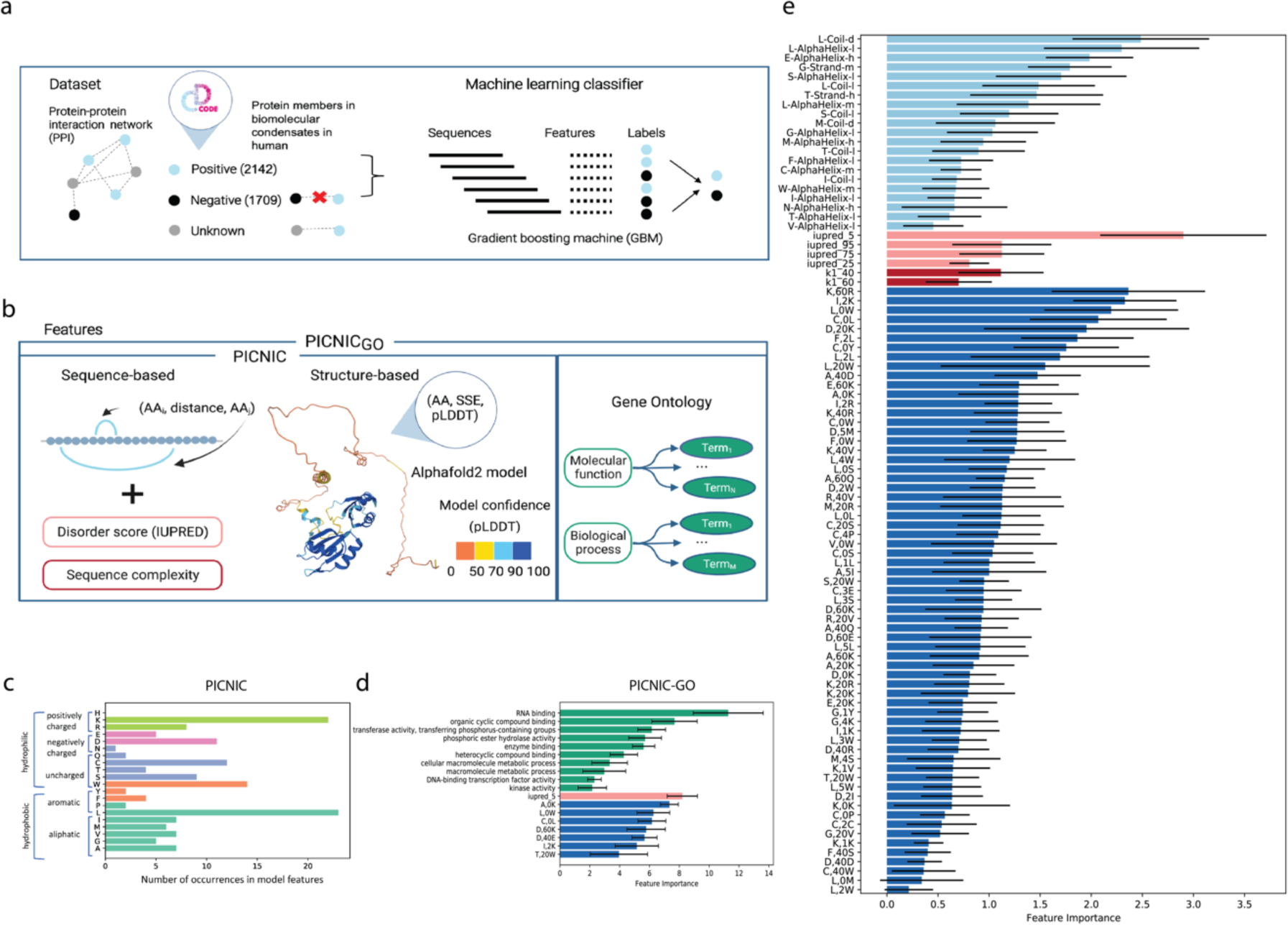
Development of PICNIC (Proteins Involved in CoNdensates In Cells) algorithm. **a)** In order to construct a training dataset, we annotated the known condensate-forming proteins from CD-CODE (positive dataset, members of biomolecular condensates) on the protein-protein interaction (PPI) network, and we excluded their first connections (proteins having interactions with condensate proteins). The remaining proteins comprised the negative dataset. Gradient boosting machine was used to distinguish two classes of proteins: members of biomolecular condensates and proteins that are not involved in any type of biomolecular condensate. **b)** Sequence, structure and function-based features of PICNIC. We developed two types of models: with (PICNIC_GO_) and without the use of gene ontology annotation features (PICNIC). Sequence-based features included sequence complexity, disorder score (IUPred), and features based on amino acid co-occurrences. Structure-based features based on Alphafold2 models included the pLDDT score, a per-residue measure of local confidence on a scale from 0 – 100 (colored on the structure). We annotated the secondary structure (SSE) based on 3D protein structures using STRIDE and all possible triads in the form (AA, SSE, pLDDT) were calculated. **c)** Amino acid occurrences in the features of PICNIC model show that Leucine and Lysine contribute the most to the model predictions. **d)** The model including GO terms (PICNIC_GO_) used three types of features, based on gene ontology (in green), disorder (orange), distance-based (violet). The most important features included RNA-binding, disorder and co-occurrences of charged and hydrophobic residues. The feature importance is consistent across different folds of cross-validation (mean values across 10 folds are shown, black lines with upper and lower border show standard deviation for each feature within cross-validation). **e)** Feature importance of PICNIC is consistent across different folds. Mean values across 10 folds are shown, black lines with upper and lower border show standard deviation for each feature within cross-validation. Features constitute three groups: based on AlphaFold2 models (in light blue), disorder (pink), complexity (dark red) and distance-based (blue).

Building the negative dataset is a complicated task as there is no publicly available resource that reports proteins that do not form condensates. Additionally, condensates may form only under specific conditions^35^. Here, we defined the negative dataset based on protein-protein interaction network (InWeb database^36^ for human proteins). We excluded all proteins that have direct connections with known condensate proteins. We reasoned that these proteins are potential condensate members that have not yet been studied. The remaining proteins comprised the negative dataset (**Figure 1a**). Of course, this procedure doesn’t guarantee the absence of condensate proteins among the negative dataset (false negatives). But exclusion of the proteins that directly interact with proteins that were reported as members of synthetic or biomolecular condensates is lowering the probability of mixing positive and negative data. Overall, our non-redundant dataset (filtered by 50% sequence identity) contained 2142 positive and 1709 negative human proteins, which were divided by 4:1 ratio into training and test datasets.

### PICNIC identifies sequence- and structure-determinants of condensate formation

We hypothesized that the ability to form condensates is encoded in the proteins’ sequence and structure, and developed a machine learning classifier called PICNIC (Proteins Involved in CoNdensates In Cell) based on sequence-distance-based and structure-based features derived from Alphafold2 models (**Figure 1b**, in total 65 sequence-distance-based and 21 structure-based features).

It has been already shown that many proteins involved in condensates harbor intrinsically disordered regions (IDRs) and low-complexity sequences. Due to their inherent flexibility, multi-valency and ability to sample multiple conformations, they are adept at a wide array of binding-related functions including macro-molecular assemblies^9,37,38^. We also tested several metrics of disorder and sequence complexity as features (Supplementary Methods). Our final model contained several features related to disorder, such as IUPred scores^33^, that have a feature importance of 0.5-3%.

Although the presence of highly disordered residues is among most important features (**Figure 1e**, pink), it is not a prerequisite for the protein to have long disordered domains to be a member of the condensate. The proportion of known condensate-forming proteins with no disordered regions in the human proteome is 21% (disordered regions < 10aa. **Figure S1**), while 33% of all human proteins have no disordered regions. For example, Human protein Guanine nucleotide exchange factor C9orf72 is a driver protein in stress granules; Speckle-type POZ protein is a driver in nuclear speckle and SPOP/DAXX body. Both proteins consist of ordered domains only that were experimentally determined by electron microscopy and X-ray crystallography, respectively (**Figure S1**, PDB ids 6LT0 ad 3HU6). Thus, both analysis of experimentally verified condensates and the selected features by our model suggest that disorder is not a necessity for condensate-forming proteins.

Along with overall sequence complexity and disorder scores of a protein, the secondary structure of individual residue types was also found to be important. We used the confidence score of the AlphaFold model prediction, the pLDDT score, that was shown to correlate with sequence disorder^39^. We represented the occurrence of an amino acid (AA) in a given secondary structure element (SSE) with a given model confidence as a triad (aa-SSE-pLDDT).

As amino acid composition bias and patterning of charges were shown to impact the ability of proteins to form condensates^30–32^, we developed features that represent short and long range co-occurrences of amino acids in the protein sequence. We represent co-occurrence of amino-acids in the protein sequence within a distance (number of amino acids in linear sequence) by triads (AA_1_, distance, AA_2_). After feature selection, the long-range distance between charged amino acids, e.g. Lysine and Arginine (K,60, R) and Aspartic acid and Lysine (D,20,K), and short-range distance of Leucine and hydrophobic amino acids (e.g. L,0,W; F,2,L; L,2,L), and the distance between Cysteine and hydrophobic amino acids were shown to be the most important features. Overall Lysine and Leucine amino acids contribute the most to the model (**Figure 1c)**.

### RNA-related functions are enriched in condensate proteins

In order to further increase to performance of our model, we integrated functional information that is already known about each protein and described as Gene Ontology (GO) terms. We developed a second classifier, called PICNIC_GO_ that combines GO terms and the previously used PICNIC features (**Figure 1b)**. After feature selection only 18 features were included in the final model, 10 new GO features and 8 features of PICNIC described in the previous section. We found, that the most significant GO term is RNA binding, which is superseding the importance of other terms and features by several orders of magnitude (**Figure 1d**).

While the GO annotation feature is biased by existing knowledge, it nevertheless validates the design of the negative dataset: it highlights RNA binding molecular function as one of the of the most important features in PICNIC_GO_ model, which is known to play crucial role in biomolecular condensate formation, as RNA molecules are significant constituents of condensates^9,40^. This feature is efficient in discriminating the two classes of proteins because the positive and negative datasets show different distributions of RNA binding annotation. In PICNIC_GO_ the following functions were marked as the most important: transferase activity, transferring phosphorus containing groups, enzyme binding, phosphoric ester hydrolase activity, organic cyclic compound binding, and heterocyclic compound binding (**Figure 1d**). The success of including functional annotation demonstrates, that for proteins with unknown cellular compartmentalization, specific set of functional descriptions can provide additional evidence for their tendency to form biomolecular condensates.

### PICNIC accurately identifies proteins involved in biomolecular condensate formation

Several data-driven predictors were developed in the last few years, that aim to predict proteins involved in LLPS from protein sequence alone or from sequence and experimental data, such as microscopy images^41^. Here, we compared the performance of PICNIC to sequence based predictors, PSAP^23^, DeePhase^42^ and two versions of PhaSePred^29^, one general model (PdPS-8fea based on 8 features) and one developed for human proteins only which uses existing experimental data, such as fluorescent microscopy images of the proteins as well as experimental data on phosphorylation sites (PdPS-10fea based on 10 features).

We compared the performance of tools on three different datasets: 1) test dataset from the recently published PhaSePred methods^29^; 2) proteins forming nuclear punctae defined by the OpenCell project^43^; 3) test dataset generated from CD-CODE^34^ (see Supplementary Methods, **Dataset S1**). Although the CD-CODE test data is not independent and was partially used by existing predictors during their training process, PICNIC has superior performance with a maximum F1-score of 0.81 **(Figure 2c)**. Not surprisingly, the models that use existing experimental information outperform the sequence-based predictors. Specifically, PICNIC_GO_ has the best performance (ROC-AUC=0.91, F1-score=0.84) followed by PdPS-10fea (ROC-AUC=0.89, F1-score=0.83).

**Figure 2.**
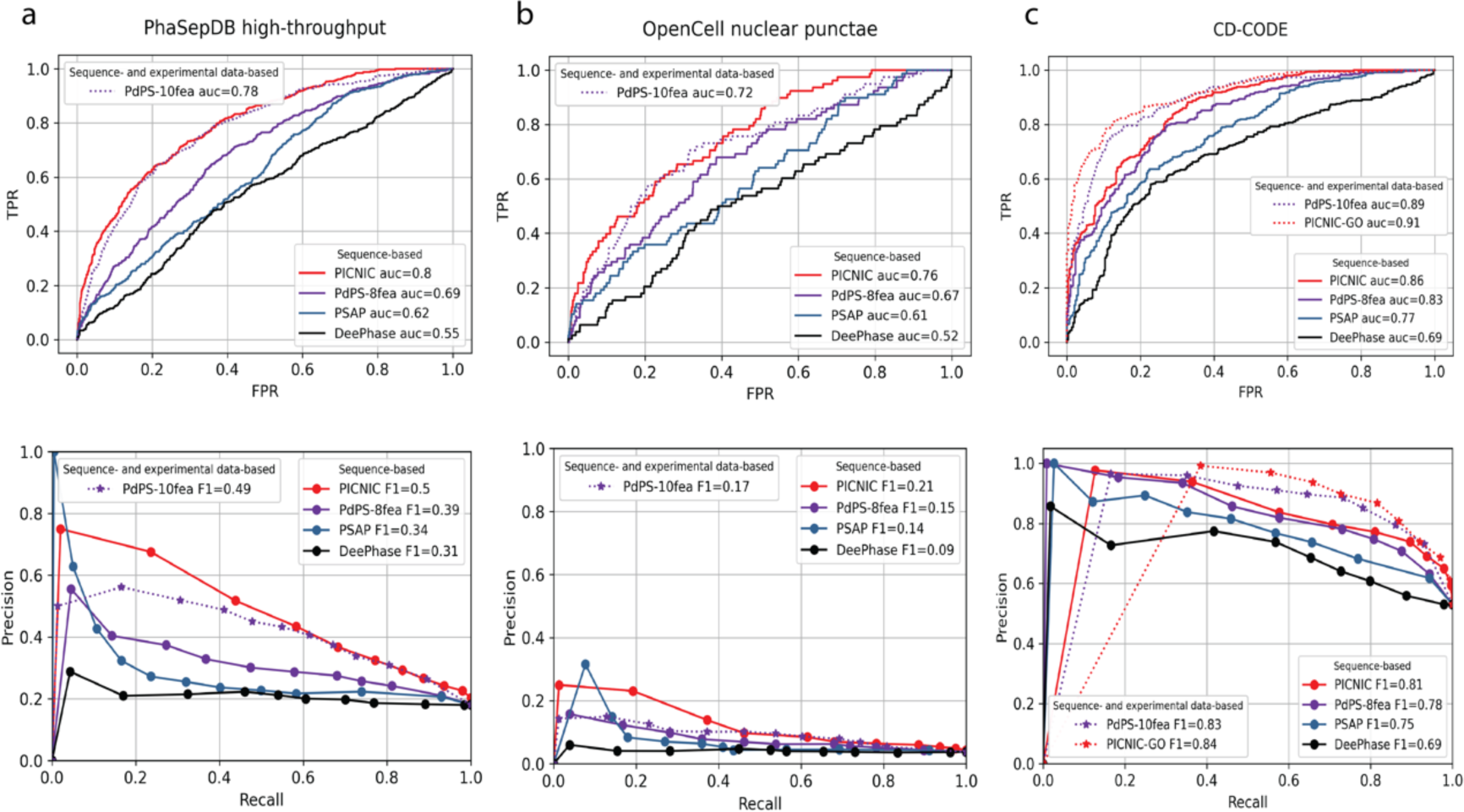
PICNIC models have the best performance in predicting condensate-forming proteins. Comparison of sequence-based predictors (lines, PICNIC, PdPS-8fea, PSAP, and DeePhase) and predictors using experimental data (dotted lines) as features to predict protein condensates. Specifically, PdPS-10-fea uses phosphorylation sites and immunofluorescent microscopy images of the proteins, PICNIC_GO_ uses GO-terms as features. **a)** Test dataset from PhaSepDB high-throughput retrieved from^29^ (441 positive and 1998 negative examples, excluding proteins that were part of the PICNIC training set), **b)** test dataset from OpenCell^43^ (78 positive and 1998 negative examples excluding proteins that were part of the PICNIC training set), **c)** test dataset from the current study based on CD-CODE^34^ (338 positive and 299 negative examples). PICNIC outperforms sequence-based predictors even on the test set that includes training data of previously published predictors, that may inflate their performance.

To further validate our model, we used microscopy images from Human Protein Atlas (HPA) where fluorescently labeled proteins were imaged and their cellular localization was determined^44^. Specifically, three types of cellular localizations were screened: nucleolus, centrosome and nuclear speckle. We filtered the list of proteins from HPA that were already in our training set that resulted in 484 proteins with known localization. Overall, PICNIC scores were higher for the proteins from HPA than for proteins without known localization (**Figure S3**). 69% of proteins mapped from HPA (with exclusion of the proteins from the training dataset) have a PICNIC score greater than 0.5, meaning that PICNIC correctly identified them as members of biomolecular condensates. It should be noted that HPA doesn’t report if a protein does not belong to given condensate (negative examples). Therefore, this dataset can be used only to check model sensitivity (recall, what fraction of true condensate forming proteins were predicted correctly), but not model precision (what fraction of positive predictions are actually true positives).

### PICNIC is robust in identifying small sequence perturbations that impact condensate formation

A real challenge of computational predictors is to be sensitive to small sequence perturbations that impact condensate formation. To test if PICNIC can distinguish similar sequences with altered condensate forming properties, we considered the synuclein family, that comprises three paralogs in human. Although they have similar sequences (**Figure 3a** and **c**, 60-70% identity) and structures as predicted by AlphaFold (**Figure 3b**), only α-and γ-synuclein form condensates *in vivo,* and only α-synuclein phase separates *in vitro*. Specifically, FITC-labeled β-synuclein, which lacks the characteristic NAC region of α-synuclein, does not phase separate at high concentrations (200 μM) and under crowding conditions (10% [weight/volume] PEG), whereas FITC-labeled α-synuclein forms condensates under the same conditions^22^. While α- and γ-synuclein can form amyloid-like fibers, β-synuclein does not^45,46^. Moreover, α-and γ-synuclein are part of biomolecular condensates: α-synuclein is reported to be the member of Synaptic vesicle pool condensate^47^, γ-synuclein is a member of the Centrosome^48^, but β-synuclein has not been found in any biomolecular condensates yet. PICNIC is the only method tested here which accurately predicts the *in vivo* condensate-forming ability of the synuclein family (**Figure 3d**).

**Figure 3.**
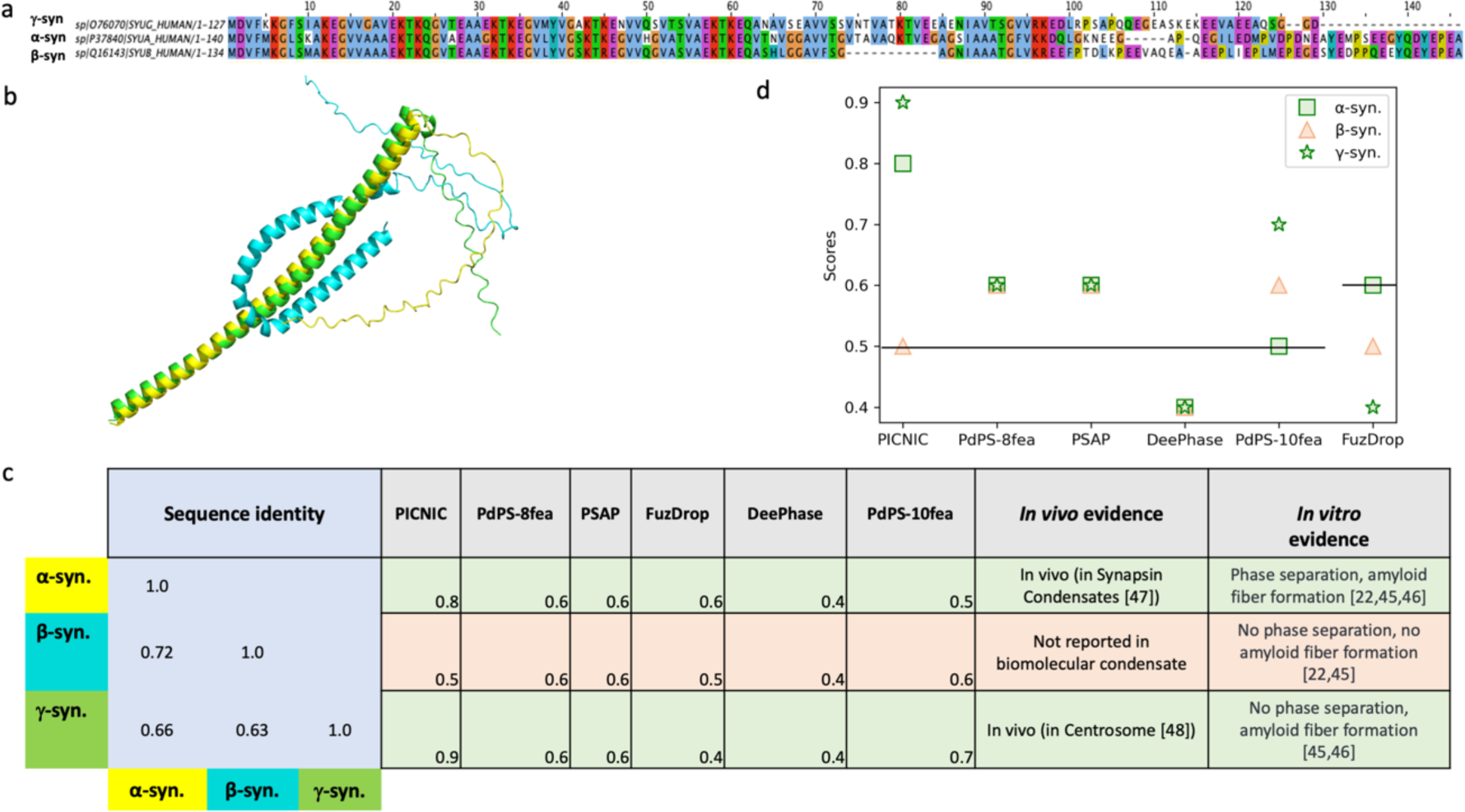
PICNIC captures the different phase separation behaviour within the synuclein protein family. **a)** The three paralogs in human share high sequence identity as depicted in the multiple sequence alignment. **b)** Structural models for α-synuclein (yellow), β-synuclein (cyan) and γ-synuclein (green), predicted by AlphaFold2 reveal that β-synuclein has a bent structure. **c)** Despite the high sequence similarity, only α- and γ-synuclein are part of biomolecular condensates, while β-synuclein has not been found in any biomolecular condensates yet and was shown not to phase separate *in vitro*. **d)** Comparison of prediction scores of different tools in identifying condensate forming (α and γ, green) and non-condensate forming paralog (β, red). PICNIC accurately predicts the condensate-forming ability of the synuclein family, and ranks β-synuclein the lowest, while other tools give equivalent scores to all paralogs or fail to identify the right trend. Vertical lines indicate the threshold used by the various methods to classify condensate-forming proteins.

Other methods either give the same score for all three paralogs and/or do not predict the correct tendency of condensate formation *in vivo*. We surmise that PICNIC is sensitive to structural rearrangements of proteins, and hypothesize that the bending of alpha-helix in β-synuclein potentially hinders the protein’s ability to form condensate.

### Interpretation of our machine learning models

To explore the generalisability of our models, we compared the most important features learnt by PICNIC vs. PICNIC_GO_. First, we calculated the most divergent gene ontology features between the dataset of known condensate-forming proteins (CD-CODE) and the dataset of whole proteomes of corresponding organisms (**Figure S4**). Further, we compared these distributions to the distributions of potential condensate proteins predicted by PICNIC. This analysis demonstrated that PICNIC can recognize the protein properties deduced as most important by PICNIC_GO_ model, as well as terms describing cellular localization which were excluded from PICNIC_GO_ features (**Figure S5**).

We also compared the two models in the opposite direction to see whether PICNIC_GO_ model can identify the sequence- and structure-based features considered as important by the PICNIC model. Interestingly, for different species different subsets of features were highlighted, but in concordance with features selected as most significant by PICNIC (**Figure S6**). Thus, PICNIC that was trained without Gene Ontology annotation can detect properties of proteins in biomolecular condensates captured by Gene Ontology terms for different species and vice versa: PICNIC_GO_ detects properties of proteins in biomolecular condensates captured by distance-based and AlphaFold-based features for different species, further validating the generalizability of the model. Overall, the propensity of a protein to be a member of biomolecular condensate seems encoded in the sequence, and machine learning models, such as PICNIC can recognize these encoded properties.

### Experimental validation of predicted condensate-forming proteins

In order to experimentally validate our model, we decided to predict the condensate localization of poorly characterized human proteins and sought to validate their condensate-forming behavior inside living cells. To do so, we chose 24 proteins (**Dataset S2**) which: i) cover diverse molecular functions spanning the entire central dogma of molecular biology and regulation of all the major cellular bio-polymers for instance, nucleic acids, proteins and chromatin (**Figure S7a**), ii) represent the average sequence length of human proteins (i.e., around 350 amino acids) by having a range of 125 to 684 amino acids (**Figure S7b**), iii) represent diverse 3D structures from ordered, alpha-helical, beta-stranded to highly disordered (**Figure 4c**) iv) are involved in genetic diseases (*AIMP1, CWC27, RP9, LMOD1*) as well as host-virus interaction (*IF2GL*). Overall, the 24 proteins, we chose for experimentally verifying and benchmarking PICNIC, represent global cellular functions and therefore are suitable to demonstrate how robust our machine learning model is in predicting condensate-forming proteins across entire proteomes.

**Figure 4.**
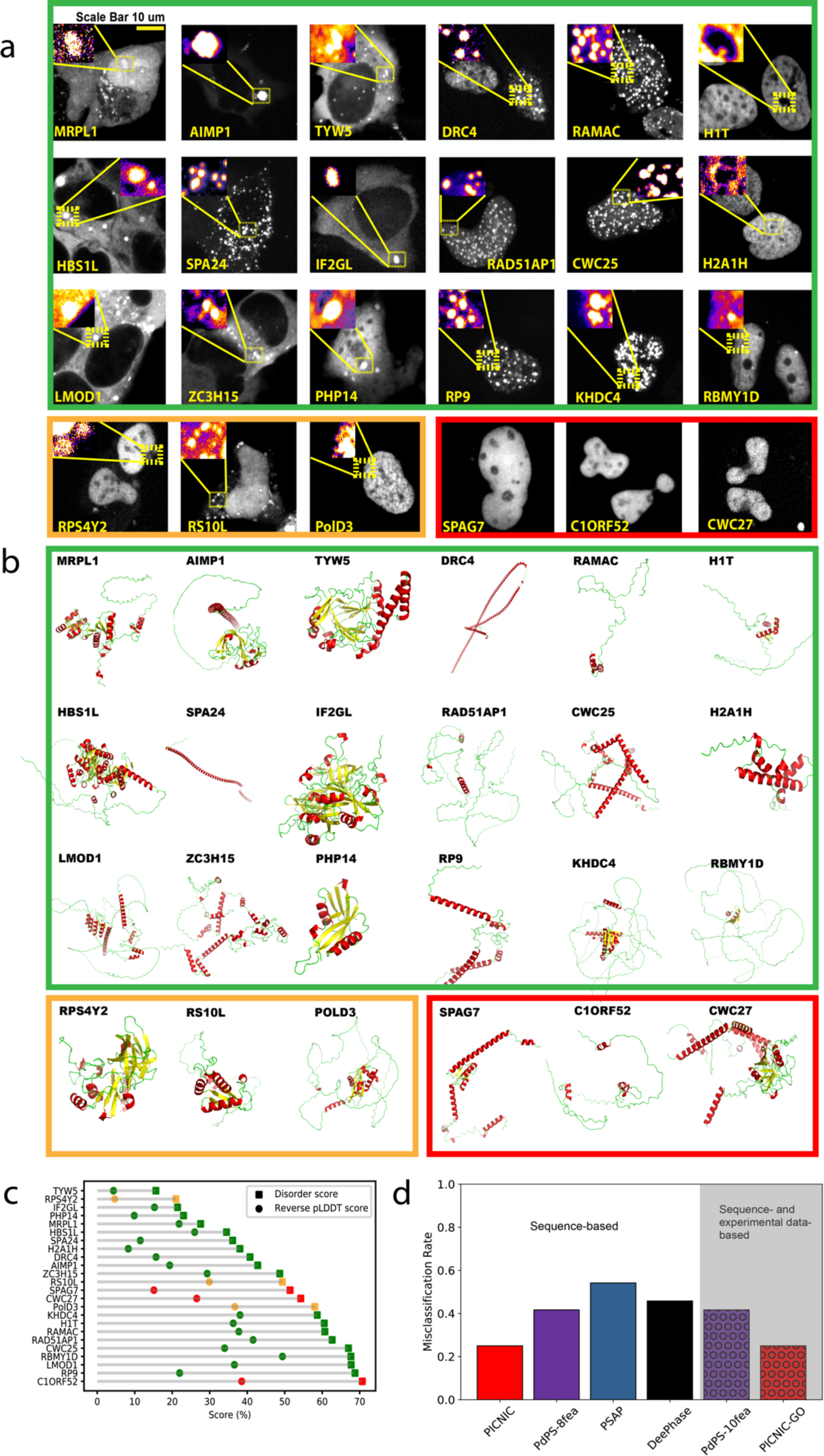
Most (18 out of 24) tested proteins form high confidence condensates *in cellulo*. **a)** Representative images of the U2OS cells expressing the tested proteins tagged with a fluorescent protein (GFP). Formation of mesoscale cellular condensates are highlighted in the inset. All images are scaled to the scale bar 10 micron (shown on upper left image). We found 21 out of the 24 tested proteins (87.5%) formed mesoscale foci without any stressors, while 3 proteins (*C1ORF52, SPAG7 and CWC27,* encircled in red) localized to the nucleoplasm without forming any detectable foci (foci were defined by exhibiting a fluorescent intensity ratio >1). Notice the presence of rim like structures in case of *H1T* and *H2A1H*. Using size, shape and the fraction of cells forming mesoscale foci as a deciding characteristic, ∼75% i.e., 18 proteins (encircled in green) form high confidence condensates, and 3 proteins (encircled in orange) form low confidence condensates (foci with longest diameter <350nm. **Figure S8**). **b)** Wide range of structural motifs covered in the test proteins; AlphaFold2 structural models of the proteins are colored according to secondary structures. Notice the wide range of structural motifs, alpha-helical (red), beta stranded (yellow) to largely disordered (green) proteins. **c)** Disorder content (computed as mean IUPred score or reverse pLDDT score (1 - pLDDT)) of the tested protein does not correlate with the ability to form condensates. **d)** Comparison of the predictions provided by sequence-based predictors (bars, PICNIC, PdPS-8fea, PSAP, and DeePhase) and predictors using experimental data (hatched bars) as features to predict protein condensates. Both PICNIC and PICNIC_GO_ exhibit the lowest misclassification rate for the tested 24 proteins.

We cloned 24 transgenes and transfected them in U2OS cells expressing fluorescently labelled proteins (see Supplementary Methods). Using fluorescent imaging, we found, that 21 out of the 24 tested proteins (87.5%) localized to mesoscale foci without any stressors, while 3 proteins (*C1ORF52, SPAG7 and CWC27,* encircled in red) localized to the nucleoplasm without forming any discernible mesoscale foci (**Figure 4a**). Foci were defined based on enrichment in fluorescent intensity, i.e., the intensity ratio inside relative to outside the foci is greater than one (**Figure S8a**). Only three proteins tested have no detectable foci, and 21 form foci (**Figure 4a**).

In order to classify the observed foci as biomolecular condensates, we aimed to define quantitative characteristics and thresholds. We measured four simple characteristics from fluorescent microscopy images (**Figure S8**): area and perimeter, informing on the size and the typical number of proteins in a foci); shape (roundness); number of foci per cell. Next, we decided on a threshold for these characteristics to aid a quantitative definition of condensates. We consider foci as condensates above the diameter of 350 nm (distance between two furthest pixels in one condensate), that is well above the diffraction limit. This would correspond to at least ∼1μm perimeter assuming a near round shape (**Figure S8c**). Using back-of-the-envelope calculations, we can consider an average protein size as 10 nm^3^, then a 1 μm^3^ compartment can contain ca. 1 million protein molecules and a 500 nm^3^ compartment can contain 100,000 protein molecules. We note, that other super resolution techniques are required to characterize the size of clusters of proteins below the diffraction limit.

### Experiments confirm 87.5% of PICNIC predictions

By applying the above definition, in our dataset, 75 %, *i.e.* 18 proteins (encircled in green) form high confidence condensates, 12.5 % i.e. 3 proteins (encircled in orange) form low confidence condensates (foci with perimeter <1 μm) while other 3 are not forming any condensates (fluorescent intensity ratio ∼ 1). We observe most condensates to be round (**Figure S8d**). The number of condensates per cell varies between 1 to 100s depending on the protein of interest.

Using co-localization experiments with known nucleolus and P-body markers, we found that 5 proteins (*R51A1, H2A1H, H1T, CWC27, MRPL1*) can localize to the nucleolus, a well-characterised liquid-like nuclear condensate (**Figure 5a**). Further, 2 proteins (*PHP14* and *HBS1L*) localize to processing bodies (P-bodies), another well-characterized cytoplasmic condensate. In addition, 8 other proteins (*RS10L, TYW5, SPA24, AIMP1, ZC3H15*, *IF2GL, LMOD1, RPS4Y2*) localized to cytoplasmic bodies and rest (*KHDC4, CWC25, PolD3, RAMAC, DRC4, RP9*) localize to the nuclear bodies (**Figure 5a**). FRAP recovery profiles for the nuclear body forming proteins revealed, that 6 out of 7 tested proteins show very fast dynamics while one (*RBMY1D*) shows no recovery at the indicated time (**Figure 5c**).

**Figure 5.**
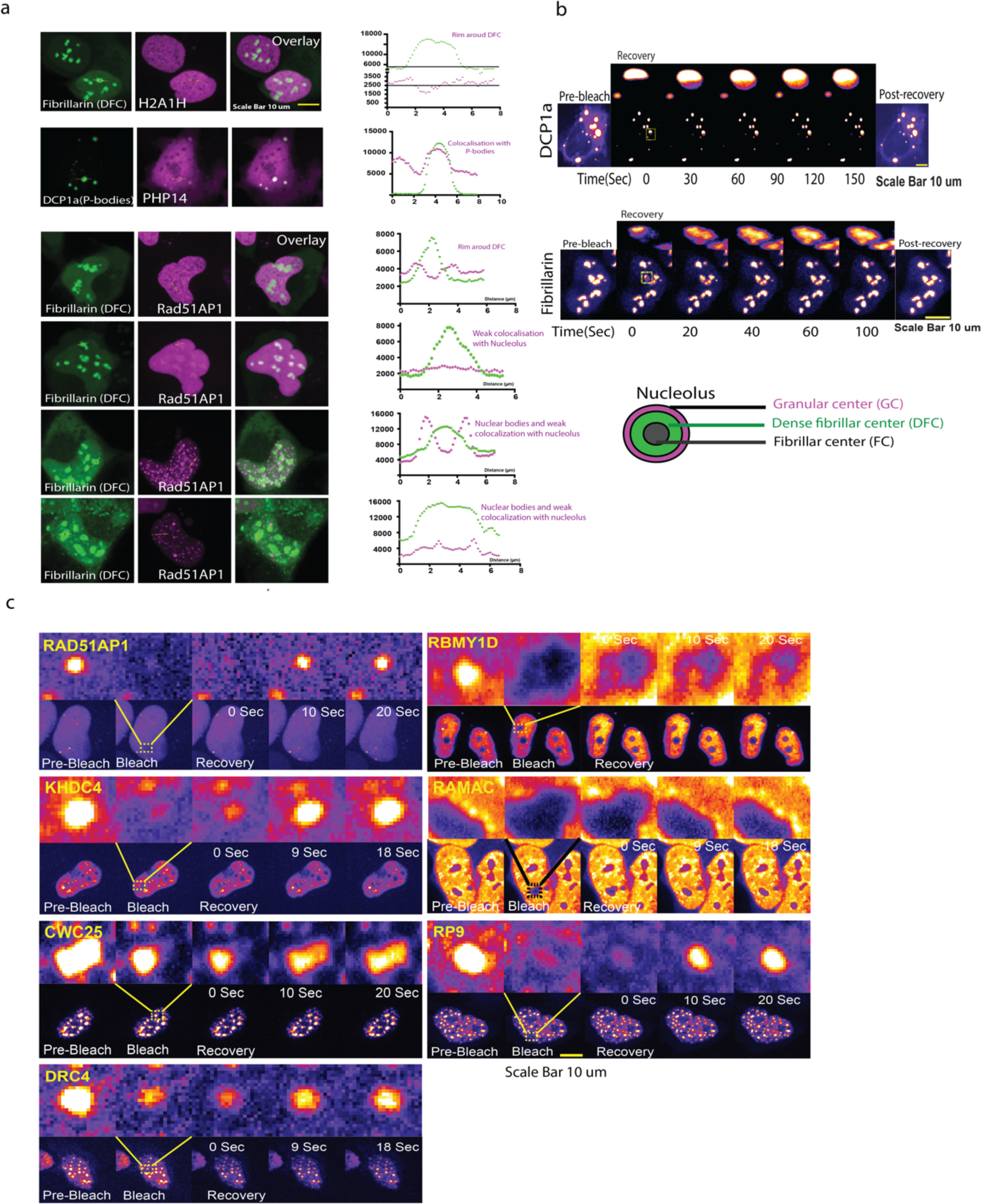
A subset of the tested proteins localizes to known condensates. **a)** Colocalization of the cellular condensate-forming proteins with well-characterized liquid like cellular condensates as can be concluded from the fluorescence intensity profiles correlation with the marker protein fluorescence profiles. While *Rad51AP1* (in purple) localizes strongly around the Dense fibrillar center (DFC, in green) forming a rim like structure, *H2A1H* (in purple) show rather weak localization as a rim around the DFC (in green) showing sub-nucleolar localization specific to the outer Granular center (GC). See the cartoon representation of the nucleolar architecture. Further, *PHP14* (purple) strongly co-localizes with the DCP1a (green) labelled processing bodies. RAD15-associated protein 1 (*Rad51AP1*) shows multi-condensate localization that varies from exclusive nuclear bodies, to nuclear bodies abutting the nucleolus, to sub-nucleolar localization (GC) as well as complete nucleolar localization suggesting an interesting regulated role for this protein’s involvement on multiple nuclear-condensates possibly in a cell-cycle stage regulated manner. All images are scaled to the scale bar 10 micron, upper right. **b)** FRAP assays showing the fast recovery dynamics consistent with the liquid like nature of the P-bodies (upper panel) and the Nucleolus (lower panel). Fibrillarin and the DCP1a FRAP recovery profile, inset highlighting the fast recovery dynamics of targeted P-body and the nucleolus. The scale bar is 10 micron each for each. **c)** FRAP recovery profiles for the nuclear body forming proteins. 6 out of 7 tested proteins show very fast dynamics while one (*RBMY1D*) shows no recovery at the indicated time. All the images with indicated whole nuclei (lower panel for each protein) are scaled to the 10-micron scale bar shown in the bottom right.

Other popular tools to predict condensate proteins would fail to make correct predictions for many of these proteins as shown by the high misclassification rates (**Figure 4d**). Overall, 87.5% of PICNIC predictions were found to be correct (misclassification rate is 25% for high confidence condensates and 12.5% if we include both high and low confidence condensates) in our experimental assays validating the model.

### Proteome-wide predictions detect no correlation of predicted condensate proteome size with disorder content and organismal complexity

In order to see if our model is generalizable, we tested its performance in identifying known condensate-forming proteins of other organisms. The CD-CODE database^34^ was screened to evaluate the fraction of proteins that were correctly identified as members of condensates by the developed model. Although PICNIC was trained on human data, it successfully predicted 72% of such proteins in mouse and 86% in Caenorhabditis elegans for example (**Figure 6a**). Thus, PICNIC model is species-independent and can be used in different organisms to assess the ability of proteins to be involved in biomolecular condensates.

**Figure 6.**
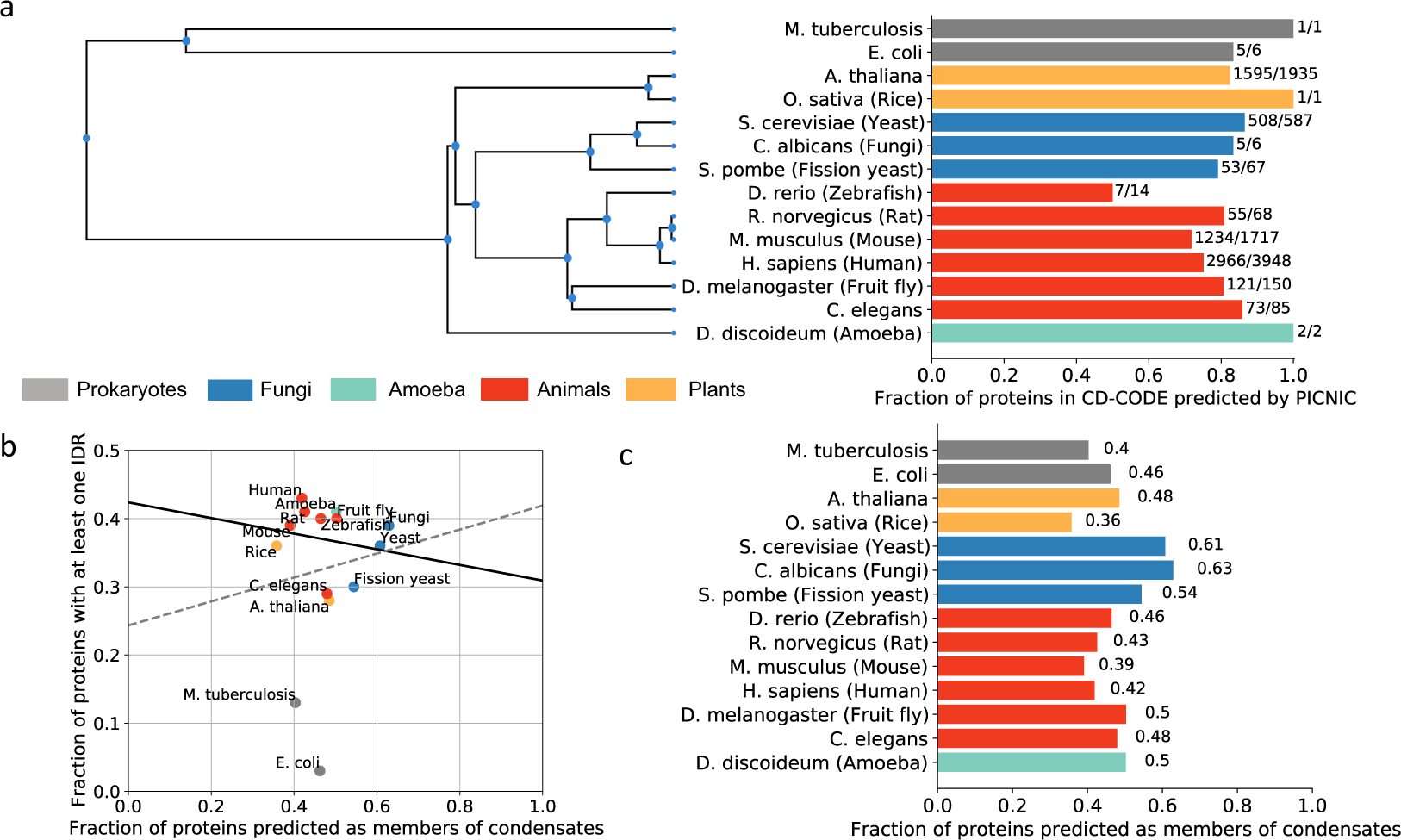
Inferring condensate proteins across the tree of life reveals no correlation with disorder content. **a) PICNIC model is species-independent.** We validated the PICNIC model on known condensate proteins from different species (defined by CD-CODE). PICNIC correctly identified 70-100% of known condensate proteins of all species tested, except for zebrafish (50%). **b) Disorder content and fraction of condensate-forming proteins of a proteome are not correlated.** Proteome-wide prediction of proteins in biomolecular condensates by PICNIC predictor (**c**) compared with the fraction of disordered proteins (proteins with at least one disordered region of >=40 residues) in proteomes shows no correlation across many organisms from bacteria to mammals (R^2^ = 0.014 for all datapoints, grey dotted line, and R^2^ = 0.033 when excluding *E. coli* and *M. tuberculosis*, black solid line).

To estimate the overall fraction of condensate-proteins, we calculated PICNIC scores for 14 different organisms across the tree of life including bacteria, plants and fungi (**Figure 6c**). We chose organisms that have already known condensate protein members that were experimentally verified in CD-CODE. We excluded organisms from further analysis where the number of known proteins is too small to compute statistics on performance: *Danio rerio* (N=14), *Dictyostelium discoideum* (N=2), *Escherichia coli* (N=6), *Mycobacterium tuberculosis* (N=1), *Oryza sativa* (N=1), *Candida albicans* (N=6). We found that the proportion of the predicted condensate-forming proteome is 40-60%, and is similar across related organisms, e.g., 42% and 39% in human and mouse, respectively (**Figure 6c**). Interestingly, while the fraction of disordered proteins increases with organismal complexity as shown before^49,50^, we found no correlation between fraction of predicted condensate proteins in a proteome and the disordered protein content (**Figure 6b**). For example, *E. coli* and *H. sapiens* have both ∼40% of their proteome predicted to be involved in biomolecular condensates **(Figure 6c**).

## Discussion

Here, we present a machine learning classifier that can learn and decode biomolecular condensate forming behaviour of proteins, that goes beyond protein structural disorder. We developed two models for the prediction of proteins involved in condensates *in vivo*, that reach precision of 77% (81% for recall) and 87% (82% for recall) for PICNIC and PICNIC_GO_, respectively at the suggested score threshold of 0.5 (**Figure 2**). The success of PICNIC models relies on several innovations: 1) novel amino acid co-occurrence features combined with protein structure-based features, that became only possible on a proteome scale since the Alphafold2 revolution^51,52^; 2) gradient boosting classifier that has an appropriate model complexity that fits the size of the training dataset; 3) improved curation and definition of the positive and negative datasets. Specifically, PICNIC models benefit from high quality positive data, that is the manually curated database of biomolecular condensates CD-CODE^34^. Additionally, we designed the negative dataset based on no *a priori* assumptions about protein disorder/structure. Previous predictors used either a i) exclusion of positive dataset from all known proteins^29,42^ or ii) proteins with 3D structures (retrieved from PDB) since phase separation is common in disordered proteins that do not have well-defined 3D structures. The simple exclusion does not provide reliable negative dataset, as it may contain many potentially phase separating proteins that have not been discovered yet. The second approach generates a biased negative dataset and is problematic because of two reasons: 1) phase separating proteins can have well-defined structure (**Figure S1), Figure 4c**), 2) such dataset is biased towards set of properties inherited by experimentally solved proteins. E.g., disorder predictors would also use 3D structures as negative dataset. To resolve this issue, here we used a protein-protein interaction network-based approach and excluded proteins that have a connection in the network with known condensate proteins (**Figure 1a**).

Gradient boosting methods (GBM) allow optimization on different loss functions which provides necessary flexibility, but more prone to overfitting. Here, the latter was compensated by choice of tree depth and providing of an evaluation dataset. GBM generally outperforms simpler models such as Random Forest or Support Vector Machine, but at the same time it doesn’t require as much data as models based on Neural Networks. Because there is not enough well-annotated data for positive and negative datasets, we chose GBM to mitigate the tradeoff between model complexity and its performance.

With the incorporation of biological knowledge about genes and proteins, PICNIC_GO_ reaches even higher performance (**Figure 2c**). Not surprisingly, it identified, along with the known impact of disordered and low complexity regions, that RNA binding seems to play crucial roles in the ability of proteins to be involved in biomolecular condensates (**Figure 1c**). Although gene ontology terms are valuable resource of information, they could introduce bias due to their nature of annotation.

We would like to emphasize that some of the other tools we benchmarked against were designed to solve a different task, namely to predict the ability of a protein to undergo phase separation or drive phase separation and *in vitro* condensate formation^22,42^. Specifically, PSAP^23^ was trained on 90 human proteins that can drive phase separation, and FuzDrop^22^ was parametrized based on 67 proteins known to self-phase separate based on in vitro experiments. Therefore, it is not expected that these tools would identify all condensate forming proteins, e.g. clients observed in vivo such as the nuclear punctae from the OpenCell project (**Figure S2b**). In contrast, PICNIC was adapted to recognize the proteins that are present in biomolecular condensates (serving a role of a driver or client) regardless of the mechanism of condensate formation. Therefore, our model does not evaluate if a protein is part of synthetic condensates, i.e., *in vitro* experiments, but rather focuses on if a protein is part of condensates in biologically relevant conditions.

PICNIC scores for clients and drivers show similar distribution (**Figure S9**). Since the same protein can behave as a driver or a client depending on the condensate identity and environment, we surmise that the ability of driving condensate formation is not solely encoded in the sequence and structure of individual proteins. This suggests, that additional input data are needed to be taken into account by the model. Studying the protein in the context of its interacting partners within a condensate may shed light on the properties of driver proteins.

While PICNIC is superior in identifying protein membership in biomolecular condensates, it has several limitations. It cannot predict which sequence features or motifs are responsible for the condensate function in a given protein. FuzDrop is the only method that can recognize specific motifs in protein sequences that promote phase separation behavior^22,53–55^.

The algorithms in this paper are designed to understand whether a protein has the potential to localize to a condensate, based on previous data. We tested 24 predicted proteins and found that 87.5% form foci *in cellulo*, and 75% form high confidence condensates based on a quantitative definition of condensates. In making this analysis, we set a cut-off above the diffraction limit. This does not mean that clusters of proteins under this limit do not form condensates, but that super-resolution techniques would be required to analyze this. It is important to state that this localization makes no claim to biological function in cells. These condensates were observed in cell lines, using GFP-tagged proteins that had varied expression levels. It is possible that some proteins will only form condensates under stress or in certain cell types at certain concentrations. In some cases, a protein for instance might not localize to a condensate under physiological conditions, but might when overexpressed in cancer. Therefore, although each protein can localize to a condensate, it remains to be sorted out by detailed experiments whether any individual protein localizes to a condensate in a specific cell type at a certain concentration and using different tags or antibodies.

We detect that structural disorder is not a prerequisite for condensate member proteins, as many of them have no IDRs both based on analysis of the CD-CODE database (**Figure S1b**) as well as 5 out of the 21 proteins identified here experimentally as condensate members have <30% disordered residues (**Figure 4c**). Thus, a wide-range of structural disorder can lead to condensate partitioning. Accordingly, we found no correlation of condensate proteome size and disorder content of an organism.

The generalizability of our model shows that the predictor learned general features (based solely on sequence information and structures that were also deduced from the sequence) across the tree of life. The provided results can shed light on evolution of biomolecular condensates across different species and by predicting condensate members beyond drivers can aid identifying potential protein targets to modulate biomolecular condensate behaviour and aid drug design^56,57^. Overall, PICNIC accurately predicts proteins involved in biomolecular condensates and provides proteome-wide perspective on proteins involved in condensate formation in different species.

## Methods

### Construction of positive and negative datasets

The positive dataset of condensate forming proteins was extracted from CD-CODE database v1.00^34^. We used all human proteins with at least 1 evidence star as a positive dataset. The negative dataset was constructed by excluding proteins that interact with known condensate forming proteins based on the InWeb v3 database^36^. After filtering the sequence for 50% sequence identity, our dataset composed of 2142 positive and 1709 negative proteins, that were divided by ratio 4:1 into training and test datasets. We divided the dataset into 20% test dataset (used only for testing the final model) and 80% working dataset, which was randomly divided into 70% training and 30% validation datasets for the 10-fold cross-validation (**Dataset S1**).

### Model features

#### Disorder score and sequence complexity

Intrinsically disordered regions (IDRs) and low-complexity regions of proteins were shown to be an important feature for predicting the ability to phase separate. To estimate protein disorder, we used the IUPred algorithm^58^, which assigns a score to each residue in the sequence. We used *k*^th^ percentile with *k* equal to [5,25,50,75,95], which is the score below which *k* percentage of residue scores fall. The 5^th^ and 95^th^ percentiles were chosen instead of minimum/maximum values to exclude the bias due to outliers.

We calculated sequence complexity in order to identify low-complexity regions (LCRs). These regions often contain repeats of single amino acids or short amino acid motifs. We calculated sequence complexity according to the definition suggested by Wootton and Federhen^59^ using two different sizes of sliding window for the protein sequence: 40 and 60. Proteins with length less than 60 residues were excluded from this analysis.

### Sequence distance-based features

We represented the co-occurrence of amino-acids (AA) in the protein sequence within certain distance by a pair of triads (AA_1_, distance_short_, AA_2_) and (AA_1_, distance_long_, AA_2_), where AA_1_, AA_2_ is one of the 20 amino acids types, distance_short_ э [1,2,3,4,5] and distance_long_ э [[0,20),[20,40),[40,60),[60,80)]. These features represent short and long range co-occurences of amino acids in the protein sequence. Short threshold distance_short_ defines the distance between two types of amino acids (e.g., 1 means neighboring residues). Long threshold, distance_long_ defines a window equal to 20, that is the distance in sequence between the two types of amino acids co-occur. The total number of tested sequence distance-based features was 1890, including both short and long distances.

### Secondary structure features based on AlphaFold models

Recent advances in Deep Learning techniques enabled *de novo* modeling of protein structures from sequences with high accuracy that is comparable to experimental methods^51^. Here, we used predicted AlphaFold2 models that were downloaded from the resource created by the EMBL Consortium (second release, Date of access: January, 2022)^52^, which contains precomputed structures for proteomes of many organisms. Alongside with atomic protein structures, AlphaFold provides the pLDDT score (predicted lDDT-Cα), that is a per-residue measure of local confidence on a scale from 0 – 100 ^60^. pLDDT scores were divided into four classes (according to DeepMind classification): [0, 50) - ‘very low’, [50,70) - low’, [70,90) –‘confident’’, [90,100] – ‘very high’. We used the STRIDE algorithm to annotate the secondary structure based on 3D protein structure ^61^. STRIDE assigns one of seven classes to each amino acid in the protein sequence: Alpha helix (H), 3-10 helix (G), PI-helix (I), Extended conformation (E), Isolated bridge (B), Turn (T), Coil (C).

Next, we calculated all possible triads in the form (AA, SSE, pLDDT), where aa belongs to one of 20 types of amino acids, SSE э [H, G, I, E, B, T, C], pLDDT э [’very low, ‘low’, ‘confident’, ‘very high’]. For longer proteins, AlphaFold2 models consist of overlapping segments of 1400 aa length. In case of discrepancy, when the same amino acid is assigned with different 3D coordinates and pLDDT score, we consolidated the predictions using the following rules: 1) We calculated all possible STRIDE predictions (with different pLDDT scores); 2) we selected the most frequent STRIDE class that had the highest pLDDT score. The total number of features based on structural information provided by Alphafold2 and STRIDE was 560.

### Gene Ontology features

Gene Ontology terms are a hierarchal dictionary of annotations describing the function of a particular gene^62,63^. They assign gene characteristics for three directions: molecular function, biological process and cellular component. Each of the direction is represented by a directed acyclic graph, where nodes represent terms (or annotations) and edges represent the relationship of subtype from descendant to ancestor node. There are tens of thousands of terms, therefore one hot encoding for all possible terms is not feasible. To decrease the number of encoded terms, we took only the most frequent terms into account. To estimate the frequency of terms, we used the Swissprot annotations of proteins (after removing redundancy by excluding sequences with more than 30% sequence identity) from Uniprot database^64^. It must be noted, that the frequency of ancestor term t_a_ also included summation of the frequencies of all descendant terms t_d_, calculated by the equation:

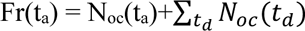 – number of occurrences of term t in the corpus.

Only terms with frequencies greater than a threshold were kept for feature calculation. Thresholds were chosen for each of the three directions separately to encompass adequate number of terms in the annotations (2500 for molecular function and biological process). We excluded the cellular component direction from the feature selection, as some gene ontology terms contain information about cellular compartments (based on Human Protein Atlas ^44^, OpenCell ^43^). We used one-hot encoding, where each protein was assigned with a vector of fixed length (equal to the number of chosen terms); 1 was assigned if the considered term in the given position (or any of their descendants) was mentioned in protein annotation, otherwise 0. The total number of features describing molecular function and biological process for the set of proteins was 1002.

### Machine learning algorithms

We developed two types of models: one including gene ontology features (PICNIC_GO_) and one without (PICNIC). Both models have the following structure: they consist of 10 Catboost classifiers with dataset for early stopping (to calculate loss function on the dataset different from training to prevent overfitting over training dataset) and fixed depth^65,66^. Catboost is a classifier based on gradient boosting machine (GBM) – a machine learning technique that gives a prediction model in the form of an ensemble of decision trees^67^.

Among other gradient boosting classifiers CatBoostClassifier from catboost library showed consistency across multiple runs (we compared LGBMClassifier from lightgbm library and XGBClassifier from xgboost library). To estimate the overall model performance across multiple runs with different parameters, we used the following metric: we chose the validation score (F1-score) of best iteration of each separate fold, and then computed the mean value across 10 folds. The model training started with all features, that is 2467 for the model without Gene Ontology features (PICNIC), and 3469 for the model with GO features, PICNIC_GO_. We selected the best features based on feature importance: at each training iteration only features with importance greater than a given threshold were selected for the next run (we took the union of features across different folds). Thus, each subsequent training iteration decreased the number of used features. We chose this feature selection approach because the feature importance did not fluctuate much for different folds (**Figure 1e**). The final models contained 18 and 92 features for PICNIC_GO_ and PICNIC, respectively. The number of features in the model with protein annotations are much lesser in comparison with another model as subset of the sequence- and structure-based features connected to phase separation properties are already directly encoded by Gene Ontology annotations.

### Cloning, cell culture and imaging

U2OS cells were transfected with plasmids encoding Human proteins tagged with iRF-670 at the N-terminus. All the 24 tested genes were codon optimized (for synthesis ease as well as to override any cellular regulation involving mRNA degradation of endogenous sequences) and synthesized by Integrated DNA Technologies (IDT) (**Dataset S3**), restriction digested using NotI-HF and AscI enzymes (NEB-R3189 and NEB-R0558) and then ligated into the pre-digested vectors using T4 DNA ligase(M0202) and transformed in *E. coli* DH5-alpha cells. Positive clones were confirmed by insert release and correctness was verified by DNA sequencing.

Cells were grown in high glucose DMEM medium (Gibco-31966021) supplemented with 10% FBS (Sigma-S0615) as well as 100 units/ml Penicillin-Streptomycin (Gibco-15140112). Following trypsinization with Trypsin-EDTA (Gibco-25300054) cells were seeded on the Ibidi 8 well imaging chamber (80826). After overnight growing the cells, cells were then transfected with plasmids encoding the 24 transgenes and for the colocalization purpose with GFP-tagged DCPa1 (a processing-body marker plasmid) and GFP-tagged Fibrillarin (a sub-nucleolar marker specific to the middle dense fibrillar region allowing the potential dissection of all 3 sub-nucleolar localization of the test proteins) using the Fugene HD transfection reagent (Promega-E2311).

Cellular fluorescent images were recorded on a spinning disk confocal microscope with FRAP capability using the 60x/1.2U-Plan-SApo, water immersion objective lens (Olympus) using 488nm and 640nm laser lines, on an Andor-iXon-897-EMCCD camera. Images, colocalization and the FRAP movies were analyzed and representative images were prepared using Fiji^68^ (**Dataset S4**).

Fluorescent microscopy images were quantified manually using Fiji and shape, size descriptors (roundness, perimeter and area) as well as fluorescence intensity were measured (**Figure S8)**. Enrichment ratio of the condensates was calculated as the ratio of the mean fluorescence of the condensate foci divided by the mean fluorescence of the background. The data was plotted using GraphPad prism software. All the images were finally prepared in Adobe Illustrator.

### Data and Software Availability

The training, validation and test datasets are available as Dataset S1. The list and properties of the 24 proteins selected for experimental validation is provided as Dataset S2. Plasmid vector maps and representative images are available as Dataset S3 and S4, shared as a public repository available at https://edmond.mpdl.mpg.de/dataset.xhtml?persistentId=doi:10.17617/3.0Y9Q8N. The PICNIC code, documentation and examples as jupyter notebooks can be found at https://git.mpi-cbg.de/tothpetroczylab/picnic. Predictions across proteomes of 14 organisms are provided as a web application https://picnic.cd-code.org.

## Acknowledgements

A.H. was funded by the ELBE postdoctoral fellowship. H.R.S thanks the Nomis foundation for financial support. A.T.-P. was funded by the MPG RGL funds. N.R. were founded by Dewpoint Therapeutics. We would like to thank the Computer Services and Scientific Computing Facilities of the MPI-CBG for their support, especially to Oscar Gonzales for supporting our HPC and HongKee Moon for developing the PICNIC web application. We thank Andrej Pozniakovski for molecular biology support.

## Author Contributions

AH, HRS, AAH and ATP designed research; AH and HRS performed research; SG and NR contributed new reagents or analytic tools; AH, HRS, SG and ATP analyzed data; AH, HRS, AAH and ATP wrote the paper.

