## Supplementary Information for "PICNIC accurately predicts condensate-forming proteins regardless of their structural disorder across organisms"

**This PDF file includes:**

Supplementary Figures S1 to S9
Supplementary Datasets S1 to S4
Supplementary References

### Supplementary Figures

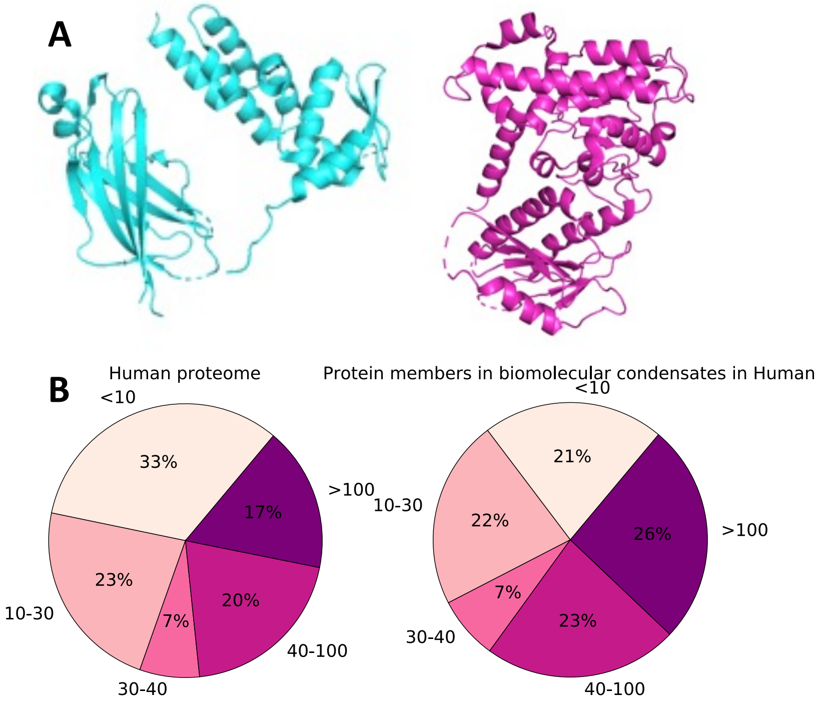


**Figure S1. Examples of driver proteins in biomolecular condensates without long disordered regions. A)** Experimentally determined structure of Speckle-type POZ protein playing role of driver in Nuclear speckle and SPOP/DAXX body condensates in human (PDB accession number 3HU6, residues 28-329 were identified with 2.70 Å resolution by X-ray crystallography) (left panel) and experimentally determined structure of protein Guanine nucleotide exchange C9orf72, which is a driver protein in Stress granules in human (PDB accession number 6LT0 chain C, residues 1-481 were identified with 3.20 Å resolution by electron microscopy) (right panel). **B)** Fraction of proteins with maximal length of disordered domain identified by IUPred (with threshold 0.5) in the human proteome (left panel) and subset of proteins in humans which are members of biomolecular condensates (right panel).

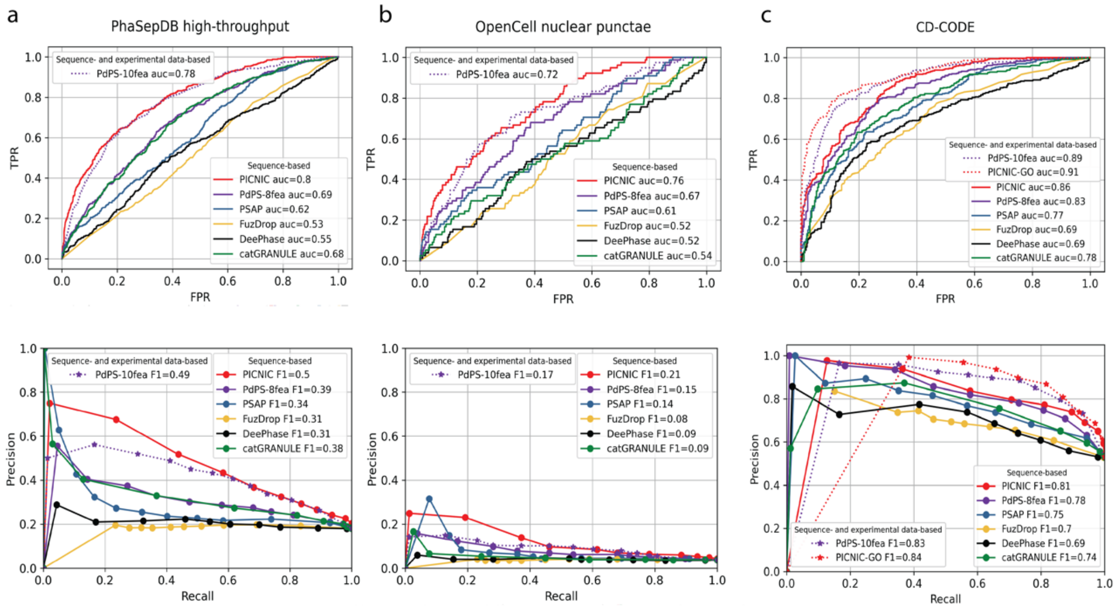


**Figure S2. PICNIC models have the best performance in predicting condensate forming proteins.** Comparison of sequence-based predictors (lines, PICNIC, PdPS-8fea[1], PSAP[2], FuzDrop[3], DeePhase[4] and catGRANULE[5]) and predictors using experimental data (dotted lines) as features to predict protein condensates. Specifically, PdPS-10-fea uses phosphorylation sites and immunofluorescent microscopy images of the proteins, $\text{PICNIC}_{\text{GO}}$ uses GO-terms as features. a) Test dataset from PhaSepDB high-throughput retrieved from[1] (441 positive and 1998 negative examples, excluding proteins that were part of the PICNIC training set), b) test dataset from OpenCell[6] (78 positive and 1998 negative examples excluding proteins that were part of the PICNIC training set), c) test dataset from the current study based on CD-CODE[7] (338 positive and 299 negative examples). PICNIC outperforms sequence-based predictors even on the test set that includes training data of previously published predictors, that may inflate their performance.

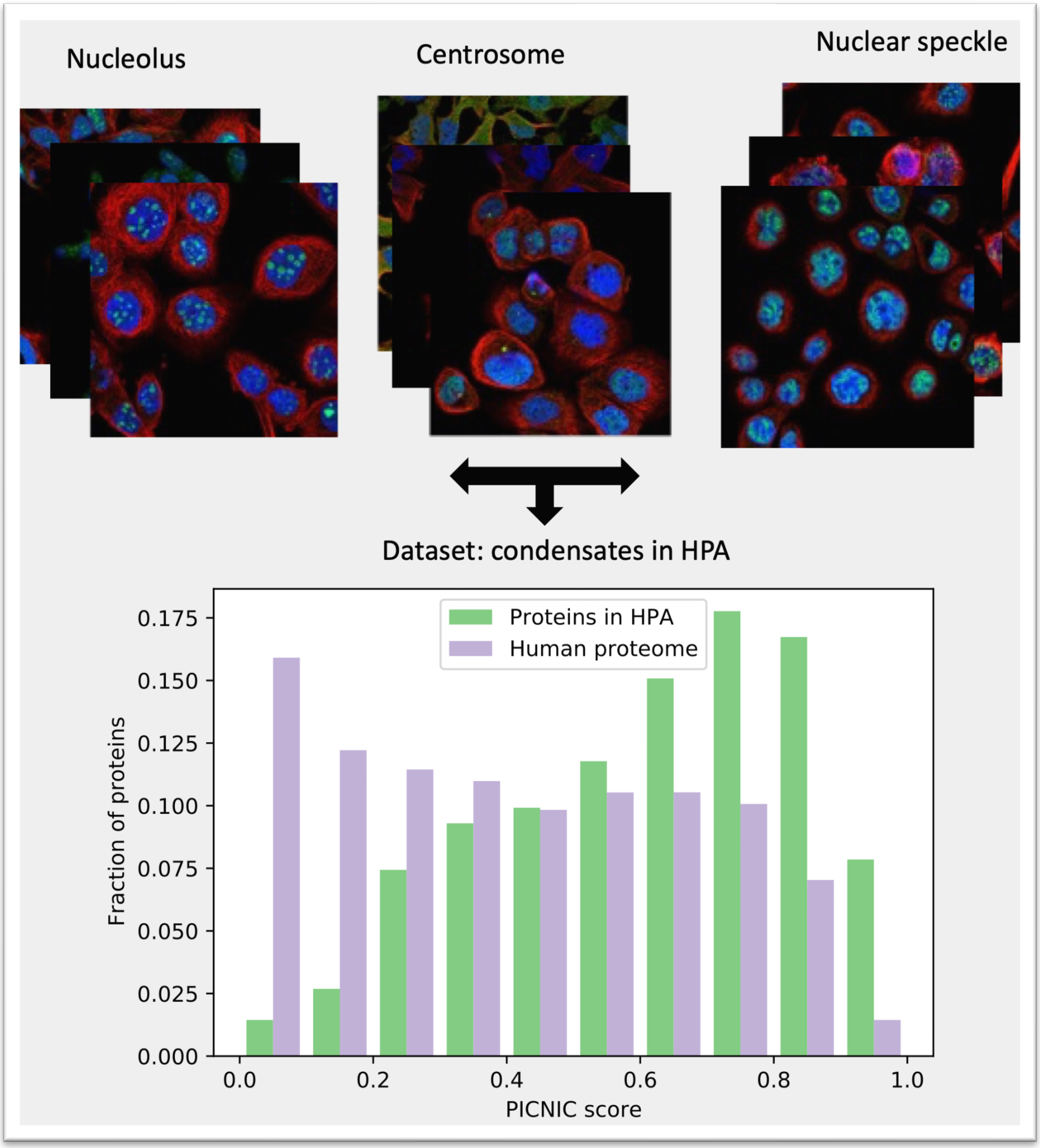


**Figure S3.** Distribution of PICNIC scores for proteins in three different condensates based on Human Protein Atlas (HPA, proteinatlas.org) (N=484, showed in green) and canonical proteins in human (N= 17361, showed in violet)[8]. Images and annotations for proteins were retrieved for three types of cellular localization: Nucleolus, Centrosome and Nuclear speckles. Only the proteins supported by experimental evidence were taken into account. All proteins from our training dataset were removed from this analysis.

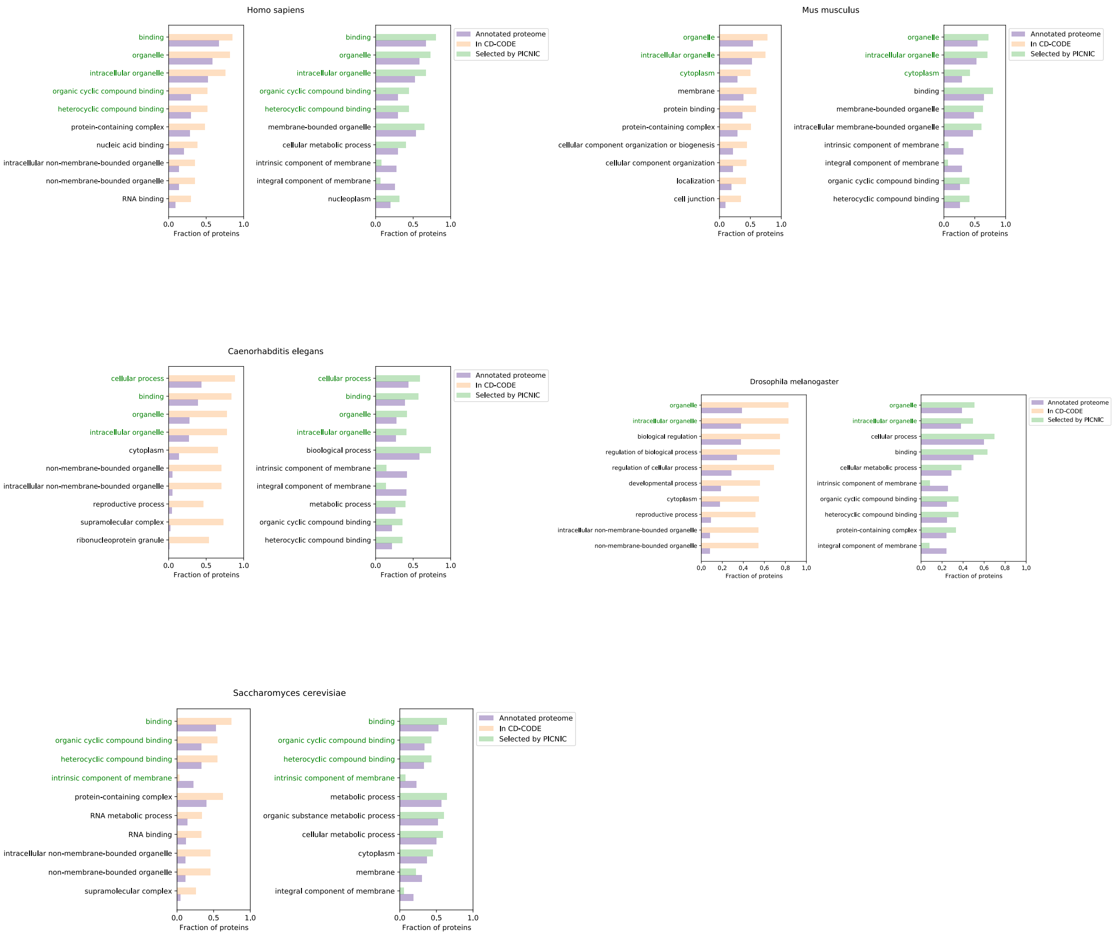


**Figure S4.** PICNIC detects properties of proteins in biomolecular condensates captured by Gene Ontology (GO) annotations for different species. Top 10 Gene Ontology features are displayed for each specie sorted by absolute value of difference in corresponding distributions (all annotated proteome vs. proteins in CD-CODE[7] on the left panel, all annotated proteome vs. proteins selected as positive by PICNIC on the right panel). Common features (with most divergent distributions between all proteins and proteins in biomolecular condensates, captured by PICNIC model) are shown in green color. Only species with sufficient number of proteins detected in biomolecular condensates (presented in CD-CODE) were chosen for the analysis.

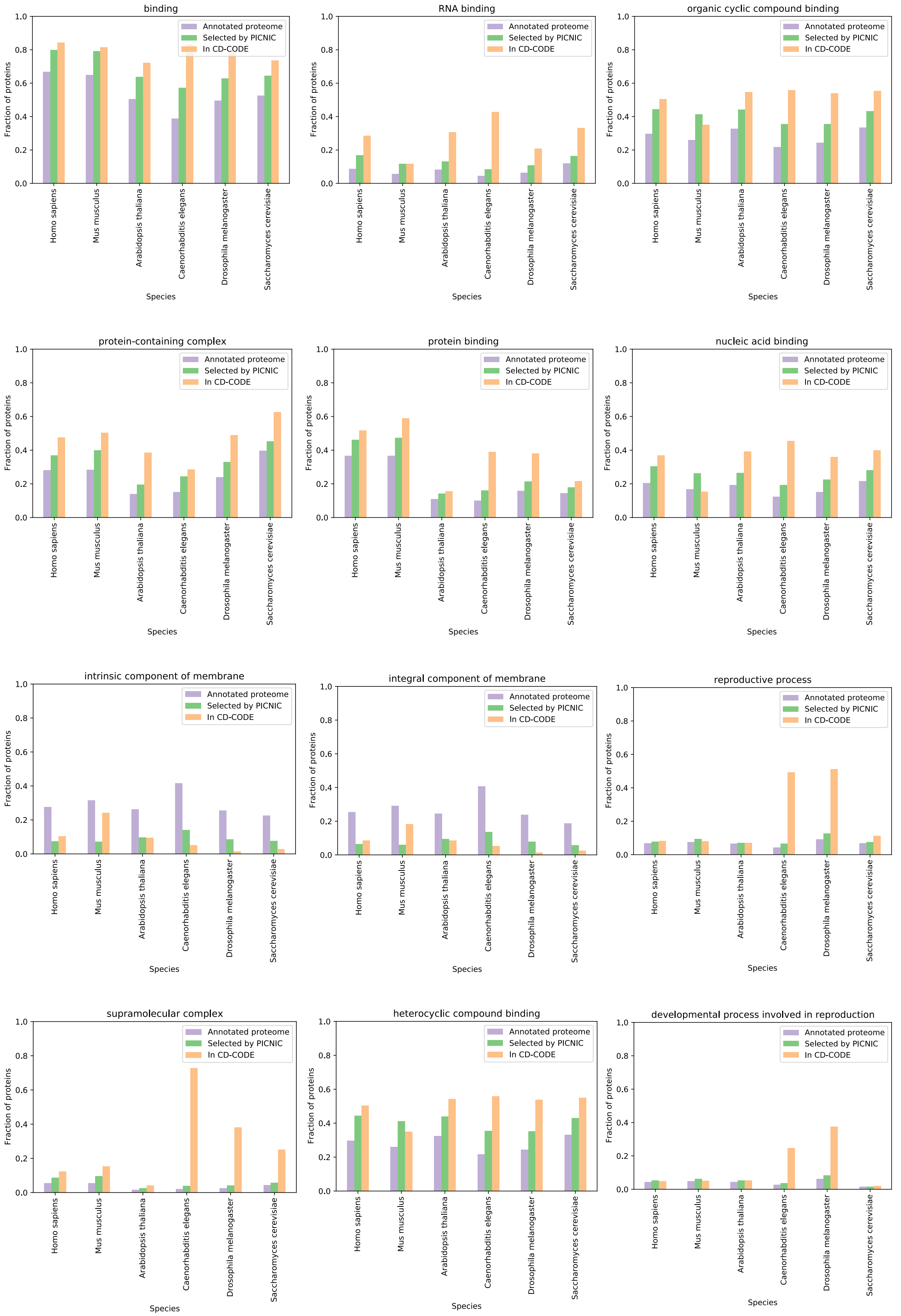
binding
RNA binding
organic cyclic compound binding
protein-containing complex
protein binding
nucleic acid binding
intrinsic component of membrane
integral component of membrane
reproductive process
supramolecular complex
heterocyclic compound binding
developmental process involved in reproduction
Annotated proteome
Selected by PICNIC
In CD-CODE
Fraction of proteins
Species
Homo sapiens
Mus musculus
Arabidopsis thaliana
Caenorhabditis elegans
Drosophila melanogaster
Saccharomyces cerevisiae

**Figure S5.** Difference in GO annotation distributions for proteins in biomolecular condensates and all proteins (grouped by GO features and compared to proteins detected by PICNIC).

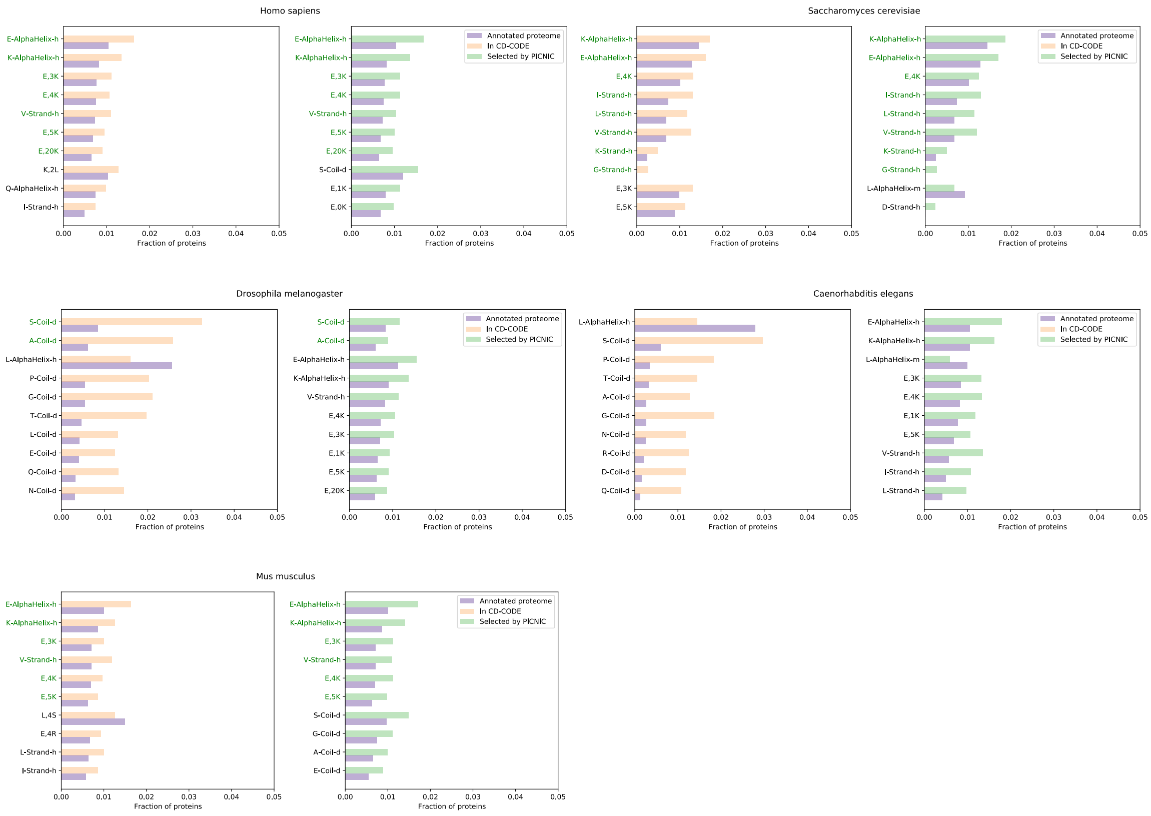


**Figure S6.** $PICNIC_{GO}$ detects properties of proteins in biomolecular condensates captured by distance-based and structure-based features for different species. Top 10 features (with exclusion of disorder and complexity score) are displayed for each specie sorted by absolute value of difference in corresponding distributions (all annotated proteome vs. proteins in CD-CODE on the left panel; all annotated proteome vs. proteins selected as positive by $PICNIC_{GO}$ on the right panel). Common features (with most divergent distributions between all proteins and proteins in biomolecular condensates, captured by $PICNIC_{GO}$ model) are shown in green color. Only species with sufficient number of proteins detected in biomolecular condensates (presented in CD-CODE) were chosen for the analysis.

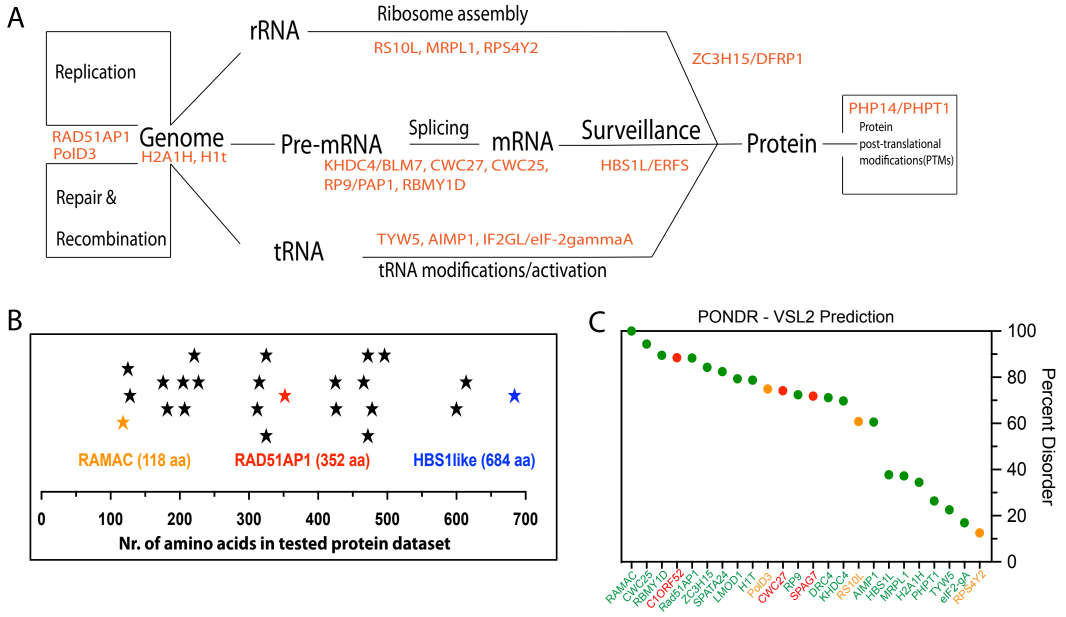


**Figure S7. Molecular functions and properties of the proteins in the experimental validation dataset.**

A) Molecular functions of the tested 24 proteins span a wide range from transcription, translation to post-translational modifications (**Dataset S2**). (B) The length of proteins varies between 125-684 amino acids. (C) There is no apparent relationship between the disorder content of proteins that form condensates (green – high confidence, yellow – low confidence) or not (red). Percent disorder was calculated using the PONDR – VSL2 webserver (pondr.com).

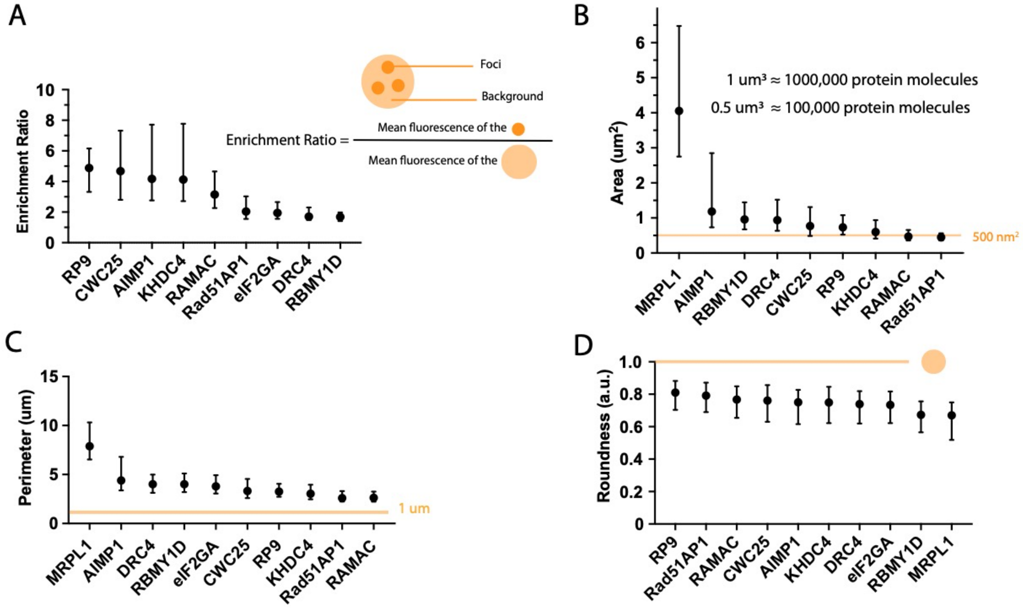


**Figure S8. Quantification of fluorescent images to define condensates.**

A) Foci were defined based on enrichment in fluorescent intensity, i.e., the intensity ratio inside relative to outside the foci is greater than one.
We measured simple characteristics from fluorescent microscopy images: area (B) and perimeter (C), informing on the size and the typical number of proteins in a foci); and shape (roundness, D). We consider foci as condensates above a diameter of 350 nm (distance between two furthest pixels in one condensate), that is well above the diffraction limit. This would correspond to ~1µm perimeter assuming a round shape. Median and interquartile ranges are shown.

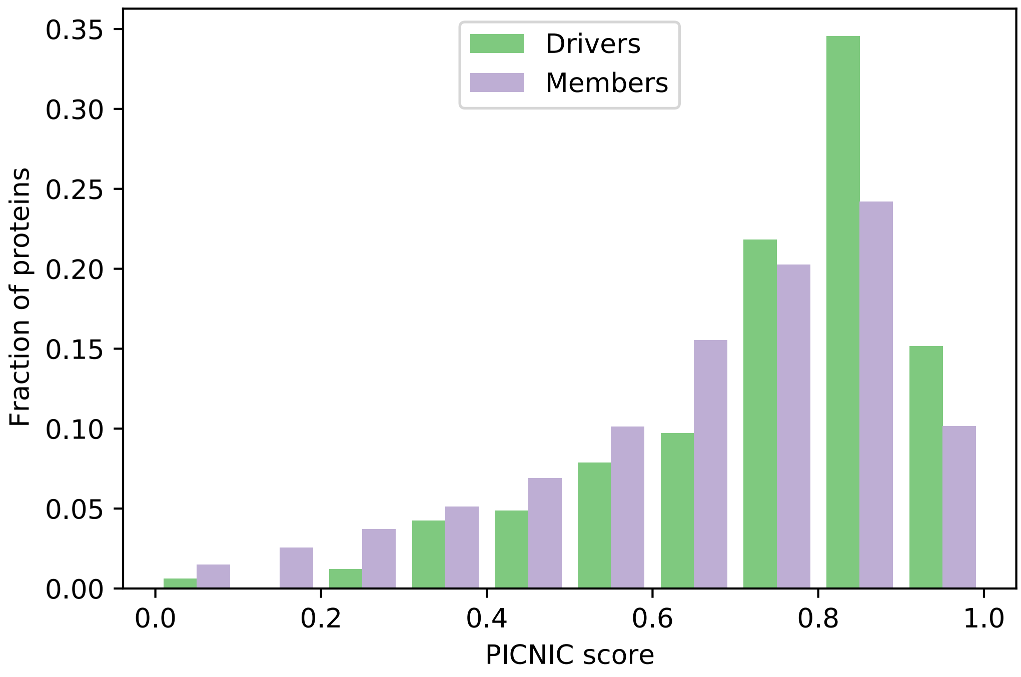


**Figure S9.** Distribution of PICNIC scores for proteins annotated as drivers (N=165 showed in green) and members (N=8074, showed in violet) in CD-CODE across 14 species.

### Supplementary Datasets

**Supplementary Dataset 1 (Dataset_S1_datasets.xlsx).** List of proteins in the training, validation and test datasets.

**Supplementary Dataset 2 (Dataset_S2_tested_proteins.xlsx).** List of proteins and their characteristics that were experimentally tested.

**Supplementary Dataset 3 (Dataset_S3_plasmid_vector_maps.zip).**

**Supplementary Dataset 4 (Dataset_S4_representative_images.zip).**
